# Octocoral microbiomes vary substantially across environmental gradients in deep waters

**DOI:** 10.1101/2023.07.05.547877

**Authors:** Samuel A. Vohsen, Santiago Herrera

**Affiliations:** Department of Biological Sciences, Lehigh University, Bethlehem, PA

**Keywords:** Mycoplasma, Oceanoplasma, Deep-sea, Mesophotic, Microbial biogeography, seascape ecology

## Abstract

Coral-associated microbiomes vary greatly between colonies and localities with functional consequences on the host. However, the full extent of variability across the ranges of most coral species remains unknown, especially in corals living in deep waters. Here we characterized the microbiomes of four octocoral species from mesophotic and deep-sea habitats in the northern Gulf of Mexico, *Muricea pendula, Swiftia exserta, Callogorgia delta,* and *Paramuricea biscaya* using 16S metabarcoding. We tested for microbiome differentiation between and within species, examining the influence of the coral’s genotype and environmental factors that vary with depth (53-2224 m) and geographic location (over 680 m). Coral microbiomes were often dominated by amplicon sequence variants whose abundances varied across hosts’ ranges including corallicolids, *Endozoicomonas*, members of the Mollicutes, and the BD1-7 clade. Coral species, depth, and geographic location significantly affected diversity, microbial community composition, and the abundance of individual microbes. Differences in host genotype, bottom temperature, and surface primary productivity could explain part of the variation associated with depth and geographic location. Altogether, this work demonstrates that the microbiomes of corals vary substantially across their ranges with potential functional consequences, identifies important ecological drivers in mesophotic and deep-sea corals, and can inform restoration efforts.

## Introduction

Shallow-water, reef-building corals are well-known for associating with microalgae in the family Symbiodinaceae. All corals, including those in the deep sea, form associations with diverse groups of bacteria [1], archaea [2, 3], unicellular eukaryotes [4], and viruses [5], collectively called the coral microbiome. Many members of the microbiome have proven important to the health of corals, either as pathogens [6] or providing sources of nutrition [7, 8].

Common coral associates include eukaryotic apicomplexans of the family Corallicolidae [9–13] and the bacterial genera *Mycoplasma* [14–19] and *Endozoicomonas* [1, 20–22]. These taxa are among the most-studied coral-associated microbes and exhibit many indications of a close relationship with their coral hosts. *Endozoicomonas* form dense aggregates in coral tissue [23, 24]. and possess capabilities relevant to the coral host such as producing vitamins [25, 26], degrading dimethylsulfoniopropionate [27], and cycling phosphorus [28]. Similarly, corallicolid apicomplexans and *Mycoplasma* are commonly found in many coral species and reside within or on coral tissue [10, 18, 29]. Most apicomplexans [30] and *Mycoplasma* spp. [31] are parasites or pathogens. However, those found in corals have not yet been associated with pathology or reduced fitness [32]. While these common coral associates have been the focus of extensive research, we have a limited understanding of the variability of coral-microbe associations.

The compositions of coral microbiomes can vary substantially between and within coral species and across geographic locations [20, 33, 34]. The genotype of the host coral underlies some of this variation [33, 35, 36], as do environmental factors such as water temperature [37], habitat type [38], and pollution levels [37, 39]. These factors can differ across the ranges of coral species, yet the scope of most studies is limited to a narrow selection of a species’ known geographic or depth range. Further, most studies have focused on shallow water corals, mainly scleractinians, and to a lesser degree on the more diverse octocorals and antipatharian black corals [34, 40, 41]. Overall, the extent of coral microbiome variability is likely underestimated.

Variations in coral microbiomes can have functional implications. In shallow-water scleractinian species, certain microbial assemblages are associated with increased thermal tolerance [42–44]. Microbiome variation could be significant for mesophotic and deep-sea corals (those living deeper than 50 meters) because they often have wider depth and geographic ranges than shallow-water species [45].

Mesophotic and deep-sea corals support diverse and abundant biological communities in Earth’s largest biome [46–50]. Most of these corals are heterotrophic because they live beyond the reach of sunlight sufficient for photosynthesis. Like shallow-water corals, mesophotic and deep-sea corals face human-induced stressors such as ocean warming, acidification, pollution, and ecosystem destruction [51–54]. In the mesophotic zone and deep sea, octocorals comprise approximately 70% of all coral species [55]. A comprehensive characterization of microbiome variation in octocorals across their geographic ranges is needed to understand the diversity and ecological roles of microbiomes in deep-water habitats..

Mesophotic and deep-sea octocorals occur worldwide along continental margins and seamounts, over a wide depth range down to at least 8,000 meters [45, 56]. Light intensity, oxygen concentration, temperature, pressure, pH, salinity, food availability, and habitat structure vary with depth and geography. All these factors combine to strongly influence the ecology and evolution of marine animals [57–60]. Thus, factors that vary with depth and geographic location likely drive microbiome variation in mesophotic and deep-sea octocorals as well.

Here we test the hypotheses that the microbiomes of mesophotic and deep-sea octocorals vary significantly across their ranges and in correlation with environmental factors and host genotype. We characterized the microbiomes of the mesophotic octocorals *Muricea pendula* and *Swiftia exserta,* and the deep-sea octocorals *Callogorgia delta* and *Paramuricea biscaya* across their known depth and geographic ranges in the northern Gulf of Mexico (**Fig. 1**). All of these coral species are dominant foundation species in their respective communities and at least three suffered mortality during the Deepwater Horizon oil spill [51, 52, 61–63] and are the focus of restoration efforts. Understanding the geographic variation of their associated microbes is essential to understanding the ecology of coral ecosystems and can also inform restoration plans.

**Figure 1:**
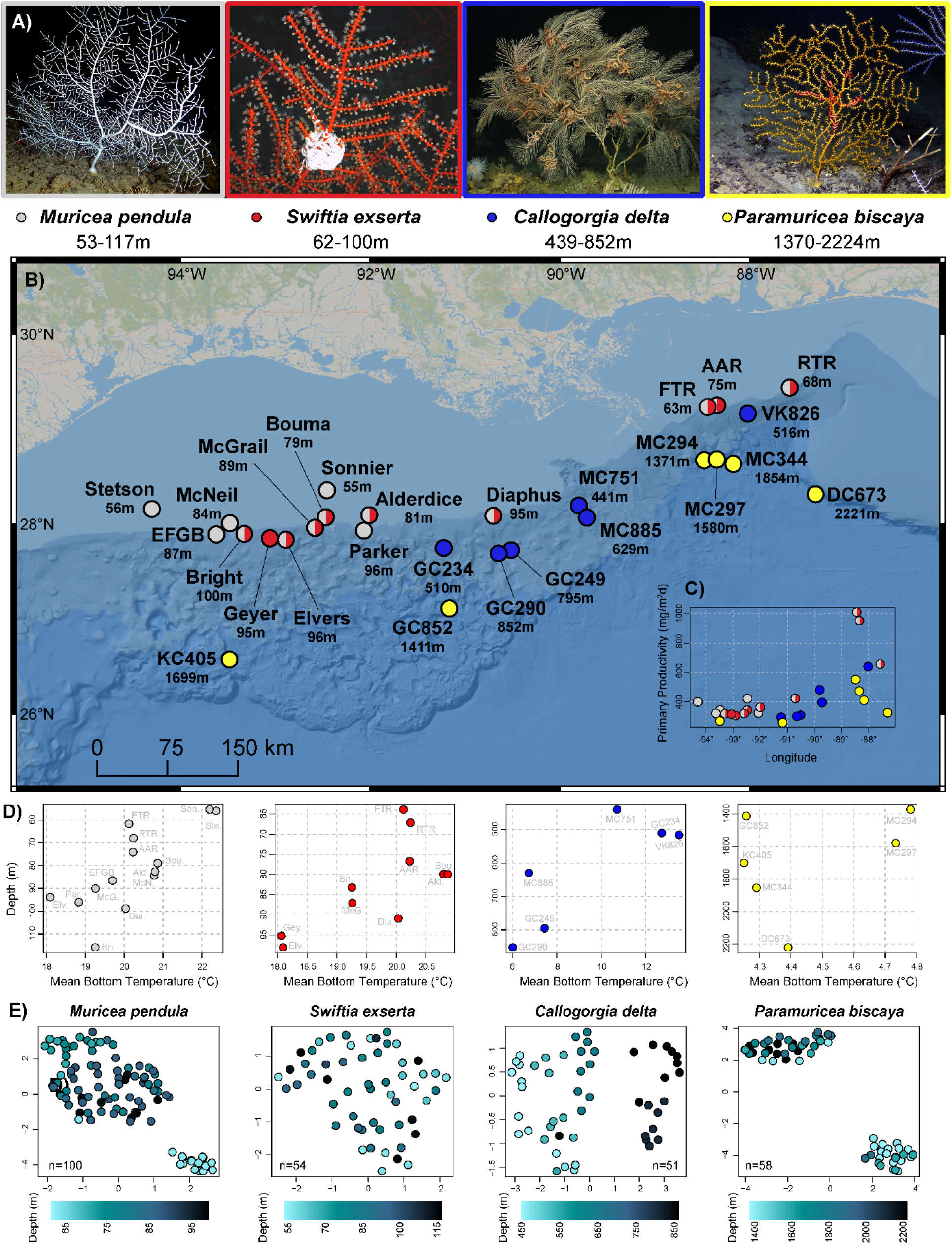
Sampling sites and their environmental and coral genotypic makeup. (A) In situ mages of the octocoral species in this study. (B) Geographic map displaying sampling locations in the Northern Gulf of Mexico (BOEM bathymetry). The mean seafloor depth of all samples from a site is reported next to each site name in meters. Sites are colored based on which coral species were sampled: light gray = *Muricea pendula,* red = *Swiftia exserta,* blue = *Callogorgia delta,* yellow = *Paramuricea biscaya.* Images of *C. delta* and *P. biscaya* courtesy of the ECOGIG consortium. (C) Primary productivity of each sampling site displayed across longitude and colored by coral species sampled. (D) Bottom seawater temperature for each sampling site displayed across depth for each coral species. (E) UMAP projections of the genetic background of coral colonies based on RADseq data. Each point represents the genotype of a coral colony and is colored by depth in meters.

## Materials and Methods

### Sample Collection

Octocorals were sampled between 2009 and 2019 from multiple sites across their ranges in the northern Gulf of Mexico using Remotely Operated Vehicles (ROVs) or the human-occupied vehicle Alvin. They were placed in temperature-insulated containers at the seafloor and processed immediately after each dive. Specimens of four species were collected including the mesophotic octocoral *Muricea pendula* (n = 140, 10 individuals per site, 14 sites spanning 34-690 km and 53-117 m in depth), the co-occurring mesophotic octocoral *Swiftia exserta* (n=100, 10 per site, 10 sites spanning 13-600 km and 62-100 m in depth), the upper-bathyal octocoral *Callogorgia delta* (n=60, 10 per site, 6 sites spanning 14-350 km and 439-852 m in depth), and lower-bathyal octocoral *Paramuricea biscaya* (n=60, 9-11 per site, 6 sites spanning 14-655 km and 1370-2224 m in depth). Details, including research cruise, time of collection, and preservation method for each sample, can be found in **Table S1**.

### DNA purification

Whole DNA was purified using a modified salting-out procedure [64]. To standardize DNA concentrations and reduce the concentration of PCR inhibitors, we measured purified DNA concentrations on a Qubit 4.0 fluorometer with a qubit Broad Range assay and then diluted with TE buffer to 2ng/µl for *Swiftia exserta*, 20ng/µL for *Muricea pendula*, 10ng/µL for *Callogorgia delta*, and 10-40ng/µL for *Paramuricea biscaya*. DNA extracts of *C. delta* and *P. biscaya* that were less concentrated than these thresholds were not diluted. See **Supplementary Methods** for details.

#### Microbial metabarcoding

The hypervariable V1-V2 regions of the bacterial 16S rRNA gene were amplified following Vohsen et al. [9] using the universal primers 27F and 355R. PCR products were sent to the University of Illinois Chicago for library preparation and sequencing [65] on an Illumina Miseq platform with 250bp paired-end sequencing.

Raw sequence data were imported into *qiime2* [66] version 2021.2 and processed following Vohsen et al. [9] to create an amplicon sequence variant (ASV) table. In brief, reads were quality filtered, joined, checked for chimeras, and ASVs were determined with deblur and classified using a naïve-bayes classifier trained on the SILVA small subunit (SSU) ribosomal RNA database Ref NR99 release 132 [67]. Details can be found in the **Supplementary Methods**.

### Coral genotyping

To investigate the influence of the host’s genotype on the microbiome, Restriction site-Associated DNA sequence (RADseq) data were obtained from a subset of coral colonies as described by Bracco et al. [58] and Galaska et al. [59]. In brief, DNA was sent to Floragenex Inc. (Eugene, OR) for RAD library preparation using the restriction enzyme, PstI, and sequencing on an Illumina HiSeq platform. Sequence reads were demultiplexed and quality-filtered using the *process_radtags* program in Stacks v2.1 [68]. Read clustering and single nucleotide polymorphism (SNP) calling were performed with DeNovoGBS v4.0.1 [69] for *C. delta* and *P. biscaya*, and with gstacks v2.1 [68] for *S. exserta* and *M. pendula* after mapping to genome assemblies for each. We excluded SNP loci that: (1) had more than 10% of missing data, (2) had a minor allele frequency smaller than 0.01, or (3) were not biallelic. We also excluded individuals that were missing data for >10% of SNP loci for *S. exserta* and *M. pendula* and >30% for *C. delta* and *P. biscaya* or were identified as clones with R package *poppr* v2.8.6 [70]. To reduce the dimensionality of the genetic data, a principal component analysis (PCA) was performed on the SNP data of each species separately (*S. exserta,* 23,869 SNPs, n = 54; *M. pendula,* 9,611 SNPs, n = 100; *C. delta*, 14,013 SNPs, n = 51 and *P. biscaya,* 7,293 SNPs, n = 58) using the R package *ade4* v1.7 [71]. The PCA axes were reduced to two dimensions using the uniform manifold approximation and projection method [72] using the R package *umap* v0.2.8. Genetic distances between individuals within species were calculated as pairwise allelic differences using the R package *poppr* v2.9.4.

### Environmental Variables

Publicly available satellite data and model-derived values for the Gulf of Mexico were obtained to evaluate the influence of environmental variables on coral microbiomes. The satellite-derived estimates of surface primary productivity (mg/m^2^d), chlorophyll concentration (mg/m^3^), inorganic suspended particulate matter concentration (g/m^3^), and the particulate back-scattering coefficient at 443nm (m^-1^) were obtained from the Copernicus-GlobColour Global Ocean Colour biogeochemical dataset. Model-derived values were obtained for the depth of the surface mixed layer (m), bottom seawater temperature (°C), bottom seawater oxygen concentration (mL/L), bottom seawater salinity, and bottom current magnitude (m/s) from the HYbrid Coordinate Ocean Model (HYCOM). Both datasets comprised monthly means from Jan 2011 to Dec 2018 at 4km grid resolution. The mean and standard deviation of monthly means for each variable were extracted at the sampling location of each coral colony. Only the mean bottom seawater temperature and surface primary productivity were retained for statistical analyses. Other variables were excluded from the analyses because they were strongly correlated, not substantially variable across sites, or were not justified by *a priori* hypotheses. See **Tables S2, S3** for further details.

### Statistical Analyses

Rarefied microbial ASV richness was calculated for each coral colony using the *rarefy* function in the R package *vegan* [73] v2.5.7 using the minimum library size (1220 reads) as the sampling depth following Hurlbert [74]. Pielou’s evenness was calculated for each individual using the function *evenness* in the R package *microbiome* v1.16.0 [75]. Microbial community richness and evenness were compared between coral species and across sites using a Kruskal-Wallis rank-sum test followed by pairwise two-tailed Wilcoxon rank-sum tests with a Bonferroni-adjusted α-value using the functions *kruskal.test* and *wilcox.test* in the base R package *stats* v4.1.2. In addition, the effects of depth and longitude on richness and evenness were evaluated for each species using a multiple linear regression model (MLR) with depth and longitude as predictors and richness or evenness as response variables.

Permutational Analysis of Variance (PERMANOVA) [76] and distance-based Redundancy Analysis (dbRDA) [77] were performed to evaluate the influence of host species, depth, geography (site and longitude), bottom temperature, and surface primary productivity on microbiome compositions. Longitude was chosen to test for regional scale patterns across geography while sampling site was used as a finer scale variable. PERMANOVA analyses were performed using the R functions *adonis2*, *dbrda*, and *anoca.cca* in the package *vegan* with Bray-Curtis dissimilarities on ASV proportions and 9,999 permutations. The results of both analyses were congruent so only the PERMANOVA results are discussed in the text. The details of each test are provided in **Tables S4 and S5**. In addition, the diversity among *Endozoicomonas* and corallicolids associated with *S. exserta* and *M. pendula* were analyzed individually using PERMANOVA to determine the influence of host species, depth, longitude, and site.

The effect of host genotype on microbiome dissimilarity was tested using Multiple Regression on distance Matrices (MRM) [78]. Bray-Curtis dissimilarities of 4^th^ root-transformed ASV proportions were used as the response variable while geographic distances, host genetic distances, and differences in depth, bottom temperature, and surface primary productivity were used as predictor variables. Geographic distances were calculated from latitude and longitude coordinates of each sample using the *distm* command in the R package *geosphere* with the Haversine function. MRM was conducted using the *MRM* command in the R package *ecodist* with 100,000 permutations.

Individual ASVs whose abundance differed along depth, longitude, and across sites were identified using Analysis of Compositions of Microbiomes with Bias Correction (ANCOM-BC) [79] using the R package *ancombc* v1.4.0. Two separate ANCOM-BC analyses were conducted on each species individually: 1) using both depth and longitude as explanatory variables and 2) a global analysis using sampling sites as groups. ASVs were excluded from the analysis if they were present in fewer than five individual corals (half the colonies sampled at a single site). The false discovery rate was controlled using the Benjamini-Hochberg method. The global analyses were repeated using each site as the reference. Only ASVs with a maximum q-value below α=0.05 in all permutations were considered significantly different. These analyses were repeated on the family level to inform functional differences.

## Results

### Microbiomes are distinct among octocoral species

The microbiomes of all four coral species were dominated by ASVs that represented more than 50% of the microbial community in at least one colony (**Fig. 2B, Table S6**). *M. pendula* had the greatest diversity of dominant ASVs, including two *Mycoplasma*, two *Marinoplasma*, an *Endozoicomonas*, an unclassified alphaproteobacterium, and a corallicolid apicomplexan. *S. exserta* hosted only two dominant ASVs: one classified as BD1-7 clade (Spongiibacteraceae) and another classified as *Mycoplasma*. Both *C. delta* and *P. biscaya* hosted three dominant ASVs. All three dominant ASVs in *C. delta* were recently described [29] as novel members of the class Mollicutes: two *Ca. Oceanoplasma* and one *Ca. Thalassoplasma*.

**Figure 2:**
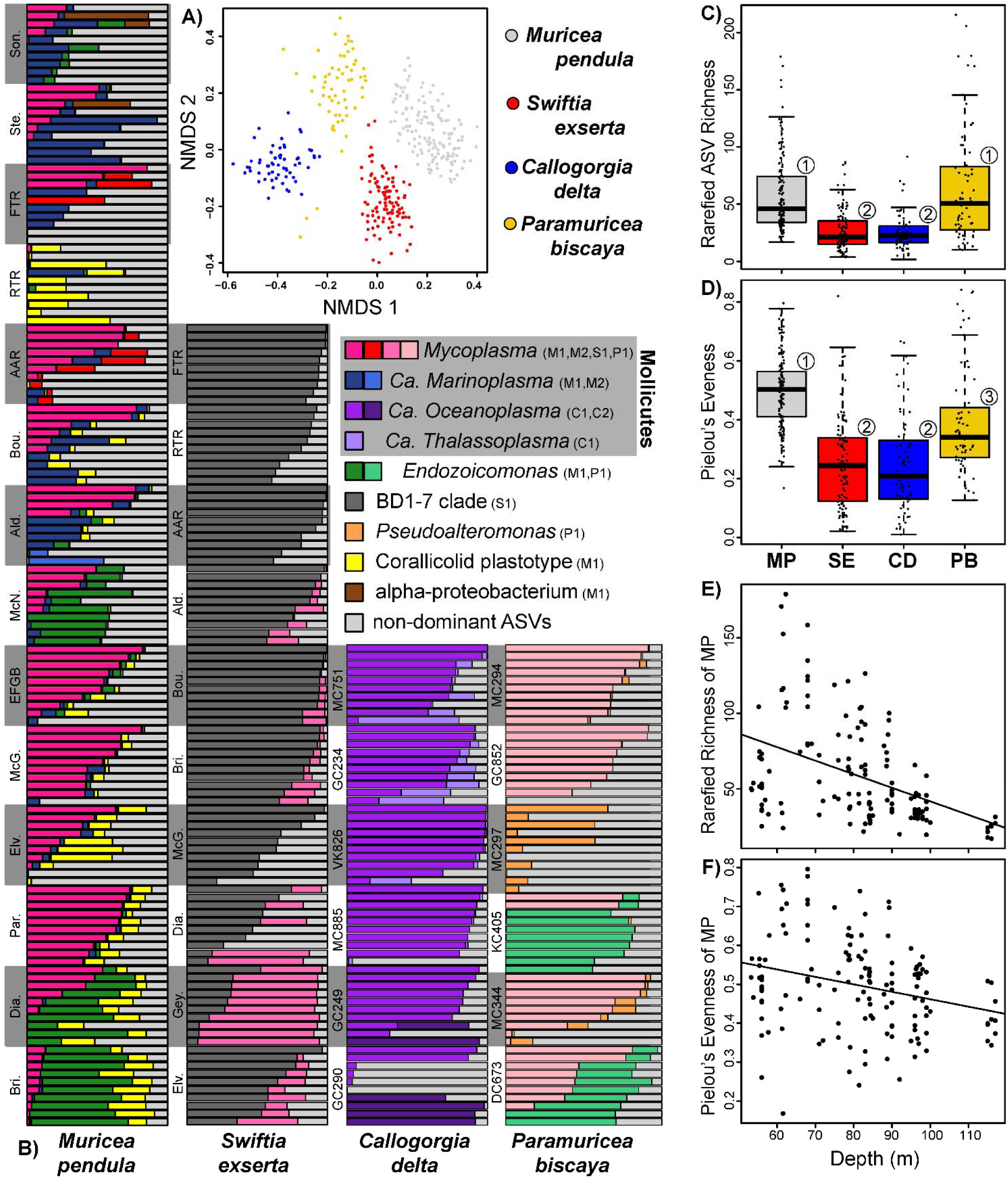
Characterization of coral microbiomes. A) Non-metric multidimensional scaling plot of microbiome compositions using Bray-Curtis dissimilarities of ASV proportions. B) Barplots of dominant ASVs in each coral species. Each horizontal bar is an individual coral colony. Colonies are grouped by site which are sorted by depth. ASVs that comprised 50% or more in any sample are designated with different colors. All other ASVs are collapsed and shown in light gray. *Ca. Oceanoplasma* and *Ca. Thalassoplasma* were recently described by Vohsen et al. (2022). C) Rarefied ASV richness and D) Pielou’s evenness between different coral species. MP = *Muricea pendula,* SE = *Swiftia exserta,* CD = *Callogorgia delta,* PB = *Paramuricea biscaya.* E) Rarefied ASV richness and F) Pielou’s evenness in *Muricea pendula* with depth. Numbers over boxplots designate significantly different groups.

The dominant ASVs in *P. biscaya* were classified as *Mycoplasma*, *Endozoicomonas,* and *Pseudoalteromonas.* Dominant ASVs belonging to the same taxon were given short identifiers (ex: M1, M2, S2) to distinguish which coral species they were associated with (first letter of coral genus) and the rank of abundance (1 being most abundant).

Each coral species had a distinct microbial community (**Fig. 2A,B; Table S4A**; PERMANOVA R^2^ 45%, p≤10^-4^). All dominant ASVs were abundant in only one coral species and were rare (<4%) in all others. The only exception was the corallicolid ASV M1 in *M. pendula* which was also abundant, but less so, in *S. exserta* along with other corallicolid ASVs.

Coral species also differed in microbiome richness and evenness (Kruskal-Wallis rank-sum test, p<2.2×10^-16^, **Table S7A**). The microbiomes of *M. pendula* and *P. biscaya* had higher rarefied ASV richness and Pielou’s evenness than *S. exserta* and *C. delta* (**Fig. 2C,D; Table S7A**; post-hoc pairwise Wilcoxon rank-sum tests, p≤5.4×10^-6^).

### Factors shaping microbiomes

The microbial communities of all coral species differed significantly across sites which accounted for 33-59% of the variation within species (**Table S4B,** PERMANOVA p≤10^-4^). The was driven by significant trends across depth (p≤10^-4^) and longitude (p≤0.0251), which simultaneously accounted for 11-17% and 3-12% of the variation, respectively, depending on coral species (**Table S4C**). The only exception was *C. delta,* whose microbial community composition was not significantly correlated with longitude. After accounting for depth and longitude, sampling site remained associated with a significant (p≤0.016) and substantial amount of additional variation (13-44%) for each coral species (**Table S4D**).

Temperature decreased with depth and was associated with a significant amount of microbiome variation in all species (**Fig. 1D; Table S4E;** PERMANOVA p≤0.0156, 5-14%) except *P. biscaya* in which it was not tested due to low variation between sites (<0.6°C). However, this did not account for the entire structure with depth which retained a significant correlation with microbiome dissimilarity after accounting for temperature (**Table S4E;** PERMANOVA p≤0.0006, 4-9%, **Table S4E**). Similarly, surface primary productivity was higher in the east and could explain a significant amount of microbiome variation in all species except *C. delta* (**Fig. 1C; Table S4F;** PERMANOVA p≤0.0154, 1-7%). After accounting for primary productivity, longitude was still associated with a significant amount of variation (**Table S4F**; PERMANOVA p≤0.0063, 4-11%,).

Microbiome dissimilarity was significantly correlated with host genetic distances while controlling for covariates in *P. biscaya* (**Table S8A**, MRM p=0.006) and weakly in *M. pendula* (**Table S8A;** MRM p=0.041). In *C. delta,* genetic distance was strongly correlated with depth (**Table S8B;** Partial Mantel p=10^-4^) and was significantly correlated with microbiome dissimilarity when ignoring depth (**Table S8C;** MRM p=0.01).

### Microbiome patterns with depth

Depth was correlated with microbiome diversity in various ways. In *Muricea pendula,* both richness (**Fig. 2E**; MLR p=6×10^-6^) and evenness (**Fig. 2F**; MLR p=0.012) decreased significantly with depth. The decrease in diversity in *M. pendula* was great enough that the average richness among colonies from the deepest sites was comparable to the richness observed in *S. exserta* and *C. delta* (**Fig. S1**). Conversely, both richness and evenness in *S. exserta* increased with depth (MLR p≤6×10^-6^), as did richness in *C. delta* (MLR p=0.038).

The relative abundances of multiple microbial taxa strongly correlated with depth including many dominant taxa. In *M. pendula*, *Endozoicomonas* ASV M1 increased in abundance with depth and so did the total abundance of all *Endozoicomonas* ASVs (**Fig. 3A**, ANCOM-BC, q=4×10^-9^, 10^-12^). Similarly, the abundance of the dominant corallicolid ASV M1 increased with depth in *M. pendula* (ANCOM-BC, q=3×10^-57^). This was also true for the total abundance of all corallicolid ASVs in *Muricea pendula* (**Fig. 3B**; ANCOM-BC q=6×10^-48^) and *Swiftia exserta* (**Fig. S2A.** ANCOM-BC q=7×10^-8^). In contrast, other dominant ASVs decreased in abundance with depth: BD1-7 ASV S1 in *S. exserta* (**Fig. 3C**; ANCOM-BC, q=5×10^-5^)*, Ca. Marinoplasma* ASV M1 in *M. pendula* (**Fig. 3D**; ANCOM-BC, q=1×10^-7^), and both *Ca. Oceanoplasma* ASV C1 and *Ca. Thalassoplasma* C1 in *C. delta* (**Fig. 2B**; q=5×10^-4^, 2×10^-52^).

**Figure 3:**
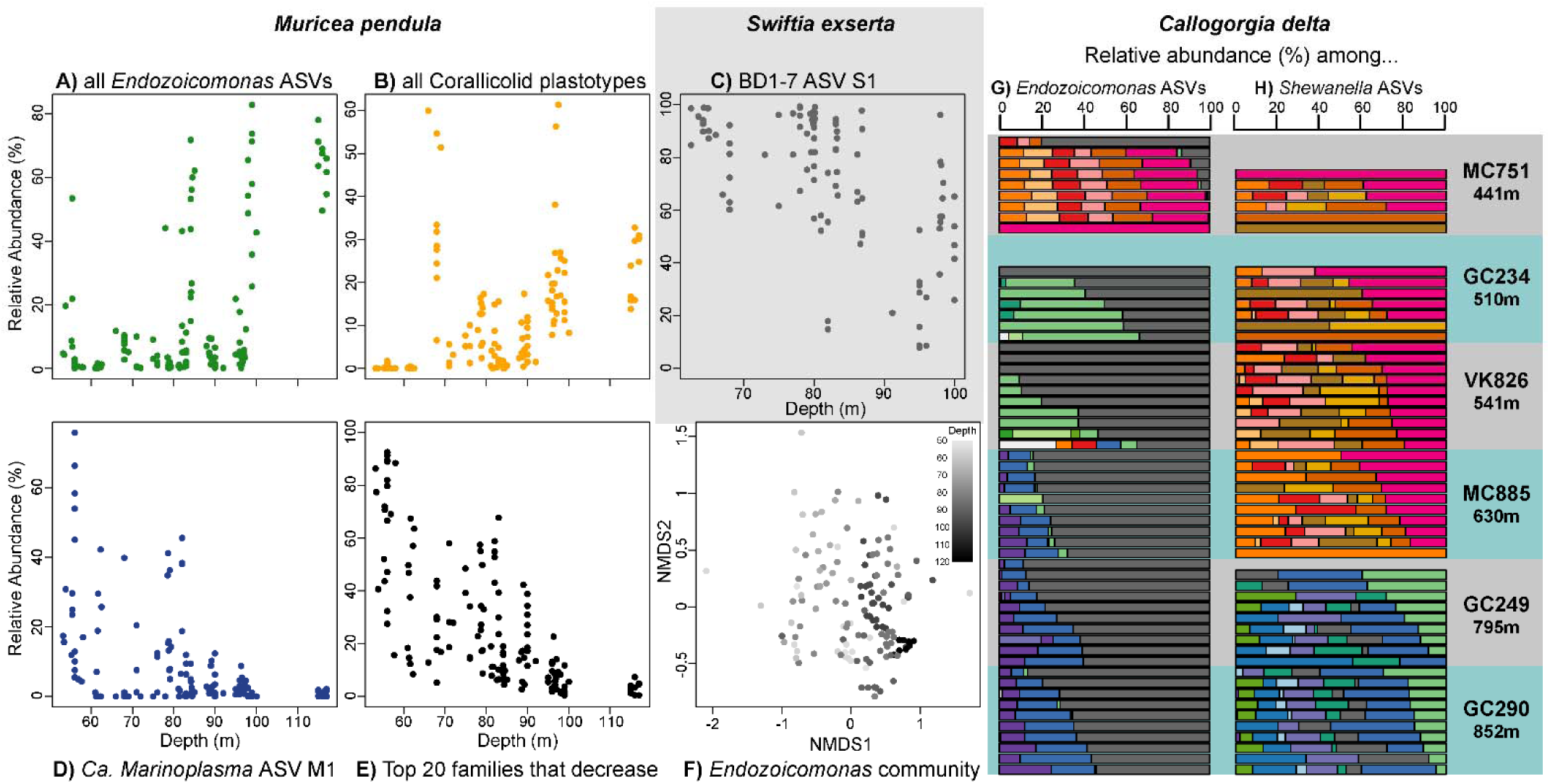
Examples of depth patterns in coral microbiome composition. The relative abundances along depth in *Muricea pendula* of A) the sum of all *Endozoicomonas* ASVs, B) the sum of all corallicolid ASVs, D) *Ca. Marinoplasma* ASV M1, E) the sum of top 20 microbial families that decreased significantly with depth. F) NMDS ordination of *Endozoicomonas* communities in *M. pendula* showing patterns with depth. C) Relative abundance of BD1-7 ASV S1 with depth in *Swiftia exserta.* The community composition of G) *Endozoicomonas* and H) *Shewanella* communities in *Callogorgia delta.* Each horizontal bar is a single coral colony whose composition reflects the relative abundances of each ASV when restricted to only *Endozoicomonas* or *Shewanella* ASVs. Colors denote different ASVs. Colonies are arranged by site in order of increasing mean depth.

Many other taxa that comprised a smaller proportion of the microbial community also differed with depth (**Fig. 3E**). *M. pendula* hosted many of these taxa and the majority decreased in abundance with depth (54/80 non-dominant ASVs and 36/42 families, ANCOM-BC q≤0.05). The most abundant of these non-dominant, depth-decreasing taxa included photosynthetic groups such as Cyanobiaceae and chloroplasts but also ASVs in the families Spirochaetaceae, Helicobacteraceae, Flavobacteriaceae, Francisellaceae, Burkholderiaceae, and unclassified bacteria that clustered with Spirochaetia and Epsilonproteobacteria. Together with the dominant *Ca. Marinoplasma* M1, these taxa averaged 70% of the microbial community among *M. pendula* colonies from the shallowest site (Sonnier Bank, 55 m) and decreased to a mean of 3% at the deepest site (Bright Bank, 116 m, **Fig. 3E**). Although many of these taxa were also detected in *S. exserta,* their abundances were either not significantly correlated with depth or increased with depth.

The composition of ASVs within microbial taxa also shifted with depth. For example, *Endozoicomonas* ASVs in *M. pendula* (**Fig. 3F, Table S4G**; PERMANOVA p≤10^-4^) and corallicolids in *S. exserta* (**Fig. S2K, Table S4G**, PERMANOVA p=0.0029). Notably, *M. pendula* and *S. exserta* each hosted a unique corallicollid ASV at the deep site Elvers Bank (96 m). Similar depth transitions were also prevalent among the associates of *C. delta*. *Ca. Oceanoplasma* ASV C1 dominated the microbial community of most *C. delta* colonies from most sites except the deepest two sites (**Fig. 2B**, GC249 and GC290). At these sites, the closely related *Ca. Oceanoplasma* ASV C2 dominated some colonies. Shifts in ASVs were also seen among *Endozoicomonas* and *Shewanella* associates of *C. delta* (**Fig. 3G,H**, ANCOM-BC by ASV, 2×10^-38^≤q≤0.002). A single *Endozoicomonas* ASV was present in most colonies from all sites (**Fig. 3G**, gray bars), while other ASVs were mostly restricted between three depth zones. More strikingly, *Shewanella* communities between colonies from two depth zones (<637M and >794m) were completely non-overlapping, failing to share a single ASV (**Fig. 3H**).

### Microbiome patterns with geography

Microbiome richness and evenness were higher in the east for both *M. pendula* and *S. exserta* (**Table S7B;** MLR p<0.0221). Multiple microbes exhibited differences in abundance across the geographic range of their host coral (**Fig. 4A-F**; ANCOM-BC, 9×10^-11^≤q≤0.05). *M. pendula* colonies from eastern sites had higher abundances of *Mycoplasma* ASV M2 and ASVs classified as *Endozoicomonas,* corallicolids, BD1-7, Spirochaetaceae, *Alteromonas, Pseudoalteromonas, Pseudomonas, Marinobacter, Ruegeria, Caedibacter,* Sphingomonadaceae, and Cyanobiaceae. The mean total relative abundance of all ASVs that were more abundant in the east ranged between 0.8% and 26%, excluding the dominant *Mycoplasma* ASV M2 and dominant corallicolid ASV M1. Some of these same ASVs and related taxa were also more abundant in *S. exserta* in the east including *Pseudomonas, Pseudoalteromonas, Alteromonas, Ruegeria, Endozoicomonas,* Cyanobiaceae, BD1-7, and Spirochaetaceae (**Supplementary File 1,** ANCOM-BC 0.0018≤q≤0.05).

**Figure 4:**
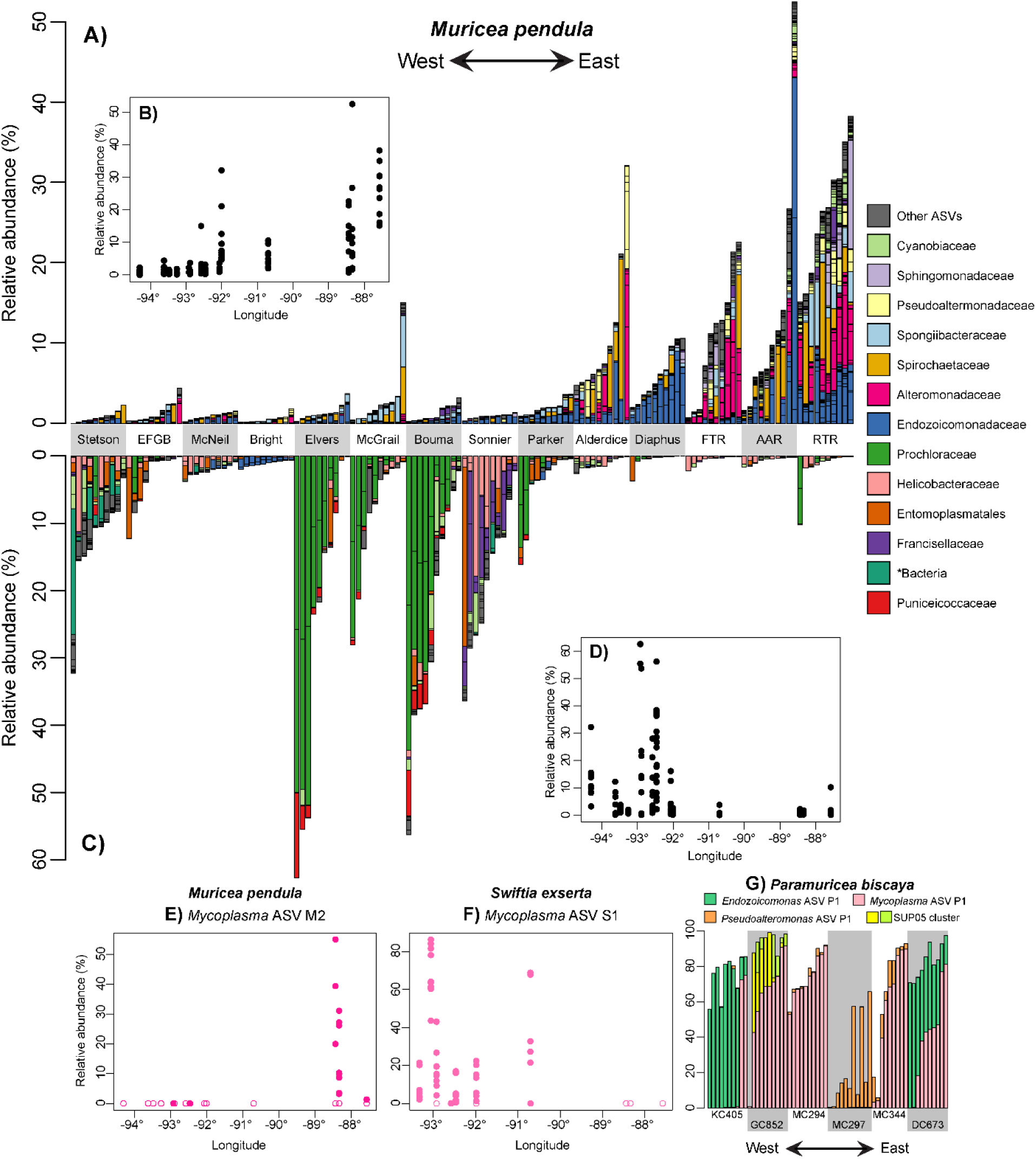
Examples of geographic patterns in coral microbiome composition. Barplots of the relative abundance of microbial ASVs from *Muricea pendula* that A) increased and B) decreased from West to East across its range in the Northern Gulf of Mexico. ASVs are colored by family or other taxonomic level detailed in the legend. Each coral colony is represented by a vertical bar, and colonies are grouped by site sorted from westernmost to easternmost starting from the left. The total relative abundances of these ASVs are shown in B) and D) plotted against longitude to better reflect relative distances between sites. The relative abundance of E) *Mycoplasma* ASV M1 in *M. pendula* and F) *Mycoplasma* ASV S1 in *S. exserta* along longitude. Samples where zero reads were detected are represented with open circles. G) Barplots showing the five most abundant ASVs that differed between sites in *P. biscaya*.

Other Cyanobiaceae, *Mycoplasma,* and *Endozoicomonas* ASVs were more abundant in *M. pendula* from western sites (**Fig. 4C,D**, ANCOM-BC, 9×10^-11^≤q≤0.03). Similarly, *Ca. Marinoplasma* and other photosynthetic taxa including several chloroplast ASVs were more abundant in the west. Notably, two ASVs classified as *Prochloron* comprised over 50% of the microbial community in some corals from adjacent sites in the west (**Fig. 4C**). Only two ASVs associated with *S. exserta* were significantly more abundant in the west including an unclassified gamma-proteobacterium (**Fig. S3G;** ANCOM-BC q=0.0018) and the dominant *Mycoplasma* ASV S1 which was completely absent from all samples from the three easternmost sites (**Fig. 4F**; q=4×10^-19^).

### Remaining Site Scale Microbiome Variation

The abundances of many microbial taxa were not correlated with depth nor longitude but still differed across sites. These comprised up to 54% of the community in *M. pendula* and 36% in *S. exserta.* Many ASVs were only detected in colonies from a few sites or even a single site. Examples include three close relatives of *Endozoicomonas* averaging a total relative abundance of 26% in *M. pendula* at McNeil bank (**Fig. S3E**) and potential parasites within Rickettsiales in all coral species. Other examples include a corallicolid ASV only detected in *C. delta* colonies from VK826, GC234, and MC885 and another corallicolid ASV only detected in colonies from GC234 (**Fig. S3H,I**).

In *P. biscaya,* the most abundant ASVs differed substantially across sites but did not correlate strongly with depth or longitude (**Fig. 4G**). For instance, SUP05 had a high abundance in colonies from GC852, *Endozoicomonas* ASV P1 was abundant at DC673 and KC405, *Pseudoalteromonas* ASV P1 was abundant at MC297, and *Mycoplasma* ASV P1 was abundant at most sites but its mean relative abundance ranged from 0.1% to 74% across sites.

## Discussion

### Differences Between Coral Species

Coral species was the strongest factor differentiating microbial communities. Species differed markedly in which microbes they associated with and also the diversity of their microbial communities. Variants of shared taxa such as *Endozoicomonas* and *Mycoplasma* were species-specific suggesting possible adaptation to different coral hosts. Other dominant microbes belonged to different phyla, such as BD1-7 in *S. exserta* and *Ca. Oceanoplasma* in *C. delta,* suggesting that the metabolic function of microbial associates may differ between coral species.

Microbiome diversity differed strongly across coral species which may reflect different ecological strategies [80]. Some corals like *M. pendula* and *P. biscaya* appear to be microbial generalists that foster more diverse and flexible microbial communities. In contrast, other corals like *S. exserta* and *C. delta* appear to have specialized communities that are less diverse and less variable [80, 81]. Alternatively, *S. exserta* and *C. delta* may have higher absolute abundances of their dominant associates reducing the richness and 16veness of their communities. Measuring absolute microbial abundances would aid the interpretation of differences in richness and evenness between species.

These comparisons between species rely on the wide sampling range of this study. *M. pendula* and *P. biscaya* had the highest microbial diversities and greater variation across sites (**Fig. S1**). However, at some sites their diversities were comparable to both *S. exserta* and *C. delta.* This demonstrates the importance of sampling widely across a host species’ range in order to fully assess microbial diversity.

### The Influence of Depth

Depth strongly structured the microbial communities of all coral species investigated. We observed differences in the composition, diversity, and abundance of specific microbial ASVs and families. The number and variety of taxon-specific patterns with depth reflect the likelihood that depth structures microbial communities through multiple mechanisms.

For instance, some variation in microbial communities may be due to temperature which decreases from >30°C at the surface to 4°C below ∼1500 m in the Gulf of Mexico. Depth and temperature were correlated with community composition in coral species living shallower than this 1500 m. Previous studies have shown that shallow-water, scleractinian corals from locations with different temperature regimes host different microbial communities [42, 82]. Specifically, the stress of high temperatures shapes corals’ microbiomes [83], especially if they bleach [84–87]. The corals in our study may experience different frequencies or intensities of thermally induced stress which may influence their microbiomes. While temperature may be the underlying cause of some of the variation with depth, it does not appear to be the only factor.

The genetic divergence of coral populations may further contribute to differentiation in microbial communities across depths. In the scleractinian *Pocillopora damicornis,* genetically distinct coral populations harbor distinct microbial communities [36]. The turnover of the *Shewanella* community in *C. delta* is consistent with its population genetic structure [57, 58]. Coral colonies from the deepest two sites were genetically distinct and hosted unique *Shewanella* communities. Thus, some of the differentiation in microbial communities across depth may be caused by genetic differentiation of the coral host.

Other patterns with depth appear to be driven by different causes. For instance, the diversity of microbial associates of *M. pendula* decreased continuously with depth and may be due to a decrease in bacterioplankton (**Fig. 2E,F**). Indeed, many bacteria from mostly free-living taxa decreased in abundance with depth in *M. pendula*. Further, light intensity may play a role since photosynthetic taxa associated with *M. pendula* predictably decreased with depth.

Altogether, depth may structure microbial communities through multiple mechanisms, including changes in free-living microbial communities, decreasing light levels, and genetic differentiation of host corals.

Many symbiotic taxa differed in abundance with depth such as *Endozoicomonas, Mycoplasma,* and corallicolids. However, the reasons for this are unclear because many of these microbe’s interactions with their coral hosts are not well characterized. Interestingly, corallicolids increased in abundance with depth in both *M. pendula* and *S. exserta,* but no depth pattern was found in corallicolids that infect *C. delta.* This is in contrast to previous studies that found a decrease in abundance with depth in the shallow-water coral, *Montanstraea annularis,* and across dozens of species spanning a depth of 2 kilometers [9, 88]. These inconsistencies may be explained if corallicolids, like their close relatives, infect intermediate hosts with differing depth distributions. Some of the closest relatives of corallicolids are found in fish [89–91], opening the possibility that corallicolids may use fish as intermediate hosts and could be limited by their depth ranges.

### Variation Across Geography

In addition to variation across depth, we found that the microbial communities of all four species differed geographically. Corals from different sites differed strongly in microbial communities in all species. Yet nearby sites showed similarities reflecting regional-scale geographic variation.

#### Regional scale geographic variation

Regional scale variation was identified in both mesophotic coral species. Both *M. pendula* and *S. exserta* had higher diversity in eastern sites where many likely free-living microbial taxa were detected at higher abundances. This may be due to the influence of the Mississippi River on bacterioplankton. There are marked longitudinal differences in surface primary productivity in the northern Gulf of Mexico, being higher in the area of influence of the Mississippi River. Microbial communities of shallow-water corals appear to be significantly influenced by surface primary productivity [92]. This is consistent with our observation that octocoral microbiomes were significantly correlated with surface primary productivity.

Symbiotic taxa also differed in abundance regionally including *Endozoicomonas,* Mollicutes, and corallicolids. This is consistent with other studies that found geographic variation in *Endozoicomonas* and *Mycoplasma* associates in shallow-water corals [15, 20]. Some of the regional patterns may result from limited dispersal since some ASVs were restricted to nearby sites like *Mycoplasma* ASV S1, which was completely absent in all *S. exserta* colonies from the three easternmost sites.

#### Site scale geographic variation

For every coral species, microbial communities differed strongly between sites after accounting for depth and geographic region. This is likely due to additional differences in local environmental conditions between sites that influence microbial community structure. For instance, the high abundance of the SUP05 cluster in *P. biscaya* at GC852 may reflect the presence of seeps nearby [93] (< 450 m). Similarly, the abundance of *Pseudoalteromonas* in *P. biscaya* at the Mississippi Canyon sites may reflect the lasting impact of the Deepwater Horizon oil spill [52, 62, 94]. Consistent deviations from depth trends at certain sites suggest that other factors play a role. For example, the total corallicolid abundance in *M. pendula* at Roughtongue Reef was higher than expected given the trend with depth (**Fig. 3B**). Conversely, corals from some subsets of sites displayed similar microbial communities despite being far apart and inhabiting different depths. For instance, *P. biscaya* from KC405 and DC673 were dominated by *Endozoicomonas* ASV P1, but it comprised no more than 0.12% in any colony from any other site. These sites are the farthest apart of all sites and do not have overlapping depths.

Interestingly, coral populations from these two sites are closely related [59] suggesting an influence of the coral host’s genotype on the presence of *Endozoicomonas*.

#### Implications for restoration efforts

Characterizations of coral microbiomes based on samples from a single site or even a few sites may miss substantial variation across a species’ range. This is important to consider in restoration plans that utilize collections from different locations. For example, a restoration plan could use this information to avoid inadvertently transmitting parasites to new sites through transplants or to avoid rearing infected and non-infected corals together in aquaria. Examples of geographic variation in potential parasites discovered in this work include corallicolids in *C. delta* at VK826, MC885, and GC234 and the unique corallicolids in *M. pendula* and *S. exserta* at Elvers Bank.

#### Conclusion

This work characterizes the microbial communities of four dominant coral species in the Gulf of Mexico and demonstrates substantial variability across their ranges. These results provide promising targets to study the interactions between corals and their associated microbes. They also reveal substantial variability associated with depth and geography that could only be uncovered through extensive sampling across the hosts’ range. The numerous patterns associated with depth demonstrate that in addition to its ecological and evolutionary effects on corals, it also strongly structures their microbial communities.

## Acknowledgements

This research was funded by the NOAA’s National Centers for Coastal Ocean Science, Competitive Research Program, the Office of Ocean Exploration and Research, and the RESTORE Science Program under awards NA18NOS4780166 and NA17NOS4510096 to Santiago Herrera at Lehigh University. Santiago Herrera was also supported by the National Academies of Sciences, Engineering, and Medicine Gulf Research Program Early-Career Fellowship under award 2000013668. We would like to thank the crews of all research vessels, ROV teams, and the other scientists involved with sampling including Destiny West and Dr. Matt Galaska. We would also like to thank Dr. Galaska for helping to generate coral genotype data.

## Data Availability Statement

16S amplicon sequence reads are available at the NCBI Sequence Read Archive (SRA) database under Bioproject number PRJNA875098. Commands used to process sequence data and R code used for statistical analyses are available on FigShare at 10.6084/m9.figshare.23656992.v1 and 10.6084/m9.figshare.21713954.v1.

