## Supplemental Information for "Octocoral microbiomes vary substantially across environmental gradients in deep waters"

**Table of Contents**


| | |
|---|---|
| **Supplemental Methods** | Pages 2-4 |
| **Supplemental Results** | Pages 4-7 |
| **References** | Pages 8 |
| **Figure S1** | Page 9 |
| **Figure S2** | Pages 10-11 |
| **Figure S3** | Page 12-13 |
| **Figure S4** | Page 13-14 |
| **Figure S5** | Page 14-15 |
| **Figure S6** | Page 15-16 |
| **Figure S7** | Page 16-17 |

### Supplemental Methods

*DNA Extractions*

DNA was extracted from 10 coral colonies from each site using a modified salting-out procedure (dx.doi.org/10.17504/protocols.io.bypypvpw). Coral tissue was digested by preparing 392uL of a cell lysis solution per sample preheated to 65°C [350µL cell lysis buffer (100mM Tris-Cl, 50mM EDTA, 1% SDS), 42µL of 0.5M EDTA, and 0.7µL β-mercaptoethanol]. About 3mm$^3$ of coral tissue was placed in the cell lysis solution, 10-30µL of 20mg/mL proteinase K was added, and samples were incubated on a shaker at 65°C for 2.5 to 4 hours. After incubation, protein and cellular debris were precipitated by adding 140µL of cold (4°C) 7.5M ammonium acetate and incubated at 4°C for 10 minutes then centrifuged at 12,000 g for 10 minutes. The supernatant was transferred to a new tube and a second round of precipitation was performed. After centrifugation, DNA was precipitated by transferring 500µL of supernatant into a new tube and adding 500µL of 100% isopropanol. Samples were inverted fifty times then centrifuged at 8,000 g for 5 minutes. DNA pellets were then washed by removing the supernatant and adding 800µL of 70% ethanol. Samples were centrifuged at 8,000 g for 1 minute and the supernatant was removed. The samples were washed a second time. DNA pellets were then air dried for 20 minutes. After drying, DNA was resuspended in TE buffer and stored at -20°C. To standardize DNA concentration and reduce the concentration of PCR inhibitors, DNA extract concentrations were measured on a Qubit 4.0 fluorometer with a qubit Broad Range assay and then diluted with TE buffer to 2ng/µl for *Swiftia exserta*, 20ng/µL for *Muricea pendula*, 10ng/µL for *Callogorgia delta*, and 10-40ng/µL for *Paramuricea biscaya*. DNA extracts of *C. delta* and *P. biscaya* that were less concentrated than these thresholds were not diluted. See **Table S1** for more details.

*PCR and Sequencing*

The hypervariable V1-V2 regions of bacterial 16S were amplified using universal bacterial primers modified with appended CS1 and CS2 linkers (underlined) 27F ACACTGACGACATGGTTCTACAAGAGTTTGATCCTGGCTCAG and 355R TACGGTAGCAGAGACTTGGTCTGCTGCCTCCCGTAGGAGT [1]. These primers were chosen because they do not amplify the mitochondrial 12S rRNA gene of corals whereas the commonly used V3+V4 primers do [2]. Twenty-five microliter polymerase chain reactions were prepared in a UV-sterilized PCR prep hood and consisted of 1X Gotaq G2 Hot Start master mix (Promega, Madison, WI), 0.25µM of each primer (Eurofins, Lancaster, PA), and 1µL of DNA extract. Thermocycler conditions consisted of an initial denaturation at 95°C for 5 minutes followed by 30 cycles of a denaturation at 95°C for 30 seconds, annealing at 51°C for 1 minute, and elongation at 72°C for 1 minute, and a final extension at 72°C for 7 minutes. Amplification was confirmed with gel electrophoresis. (3uL PCR product, 1% TE agarose gel with 1X gelred, 1X TBE buffer, 110V for 30 minutes). PCR products were sent to University of Illinois Chicago for library preparation and sequencing [3] on an Illumina Miseq platform with 250bp paired end sequencing.

*Data Processing*

Raw sequence data was imported into qiime2 [4] ver2021.2 as fastq files. Read pairs were joined using vsearch [5] with default parameters and quality filtered using quality-filter q-score [6] with default parameters. Chimeras were removed and amplicon sequence variants (ASVs) were determined using deblur [7] trimming sequences to 300bp. 1,953,131 total reads remained after filtering (1,220 – 13,712, median = 4,988). ASVs were classified using the SILVA 132 SSUref nr99 following Bokulich et al. [8] by first extracting the V1-V2 region of these reference sequences and truncating to 400bp, creating a Naïve Bayes classifier using the reference sequences, then classifying ASVs using scikit-learn [9]. A phylogenetic tree was constructed to infer the taxonomy of unclassified ASVs. ASVs were aligned with MAFFT ver7 [10] and masked by only retaining columns in which at least one non-gap character was present in at least 40% of sequences as in Lane [11]. A midpoint-rooted phylogenetic tree was constructed using FastTree 2 [12] which generates approximate maximum-likelihood trees for very large alignments (1,000s of sequences).

*Additional Statistics*

The effect of depth on genetic distance was tested while controlling for geographic distance using partial mantel tests with the mantel.partial command in the R package *vegan* v2.6-4. Bray-Curtis dissimilarities on ASV proportions, pearson's correlation, and 9,999 free permutations were used.

### Supplemental Results

*Core ASVs*

To further characterize microbial communities, core ASVs were identified that were present in at least 50% of colonies for each species (**Table S6**). Core ASVs excluding those that were also reported as dominant ASVs are as follows. In *M. pendula,* these ASVs were classified as *Helicobacter, Achromobacter, Endozoicomonas, Spirochaeta 2, Alteromonas, Fangia,* BD1-7 clade, two additional corallicolid plastotypes, Spirochaetia, and Flavobacteriaceae. Those is *S. exserta* include an additional member of the BD1-7 clade, the *Achromobacter* and two corallicolid plastotypes that were also core in *M. pendula, Sporolactobacillus, Stenotrophomonas, Serratia, Delftia,* and novel members of the Rhodobacteraceae, Terasakiellaceae, and Mollicutes. In *C. delta,* core ASVs included *Endozoicomonas,* members of the SUP05 cluster, and other novel members of the Mollicutes, Spirochaetia, Gracilibacteria (3), Thiobarbaceae (2), and Thioglobaceae. Finally, core ASVs in *P. biscaya* include another

*Pseudoalteromonas*, the *Alteromonas* that was core in *M. pendula*, the *Stenotrophomonas* that was core in *S. exserta*, the *Achromobacter* that was core in both *M. pendula* and *S. exserta*, and the most abundant novel Mollicute in *C. delta*.

*Effects of Sampling Year and Preservation Method*

Samples were collected across multiple years including 2009, 2010, 2017, 2018, and 2019. Most samples comprised snap frozen tissue however some were preserved in ethanol and RNAlater. *Muricea pendula* and *Swiftia exserta* samples were collected in 2017, 2018, and 2019. All were snap frozen tissue except the *Muricea pendula* samples from Stetson Bank which were all preserved in ethanol. *Callogorgia delta* samples were all collected in 2017 except those from MC885 (all collected in 2010) and VK826 (three in 2009 and seven in 2010). All samples were frozen tissue except those from MC885 which were all preserved in RNAlater. All *Paramuricea biscaya* samples were collected in 2017 except one colony which was collected in 2009 and not used in the final analyses. Most samples were frozen tissue except the unused sample from 2009 which was preserved in RNAlater and seven in ethanol (four from MC297, one from MC294, and five from MC344). For five colonies preserved in ethanol, DNA was also extracted from frozen tissue and sequenced. The samples preserved in ethanol were used in analyses because the extracts from frozen tissue had low concentrations and did not amplify as strongly.

To determine whether any results were due to preservation method or sampling year we repeated analyses while incorporating preservation method and/or sampling year. Preservation method was not considered for *Swiftia exserta* because all samples were frozen tissue and sampling year was not considered for *Paramuricea biscaya* because all samples in the final analyses were collected in 2017. In all analyses, a conservative approach was taken to prioritize preservation method and sampling year.

Linear models of richness and evenness were repeated using preservation method and sampling year as predictor variables. Most significant results remained except richness was no longer significant with depth in *C. delta* and evenness was no longer significant along longitude in either *M. pendula* or *S. exserta*.

All PERMANOVA analyses were repeated while incorporating both preservation method and sampling year. In analyses where terms were added sequentially, preservation method and sampling year were added first. All significant results were consistent except sampling site did not explain a significant amount of variation in *C. delta* after controlling for both depth and longitude (**Table S9**). The amount of variance explained by each variable was expectedly less due to the variance that could be explained by preservation method and sampling year.

ANCOM-BC analyses were repeated that identified ASVs that differed along depth and longitude while incorporating both preservation method and sampling year as predictor variables (**Supplementary File 1**). Most significant ASVs remained significant for all species. In general, the ASVs with the largest effect sizes and lowest q-values were consistently significant.

**Table A**. Number of ASVs that were significant across depth and longitude while using preservation method and sampling year compared to ignoring those covariates.

| | *Muricea pendula* | *Swiftia exserta* | *Callogorgia delta* | *Paramuricea biscaya* |
|---|---|---|---|---|
| Depth | 58/84 | 14/14 | 45/45 | 72/75 |
| Longitude | 159/164 | 22/22 | 49/77 | 179/195 |

Additionally, we analyzed a similar dataset including *C. delta* colonies from the same sites but collected in 2015, 2016, and 2017 using both frozen tissue and tissue preserved in ethanol. PERMANOVA analyses were repeated. Only significant effects were compared and the magnitude of $R^2$ values were ignored since sampling across sites was not uniform in this dataset. All significant effects were replicated (**Table S10**). Microbiome dissimilarities differed significantly between sites and were correlated with depth and longitude. Site explained a significant amount of variation after controlling for depth and longitude. Temperature could explain some of the pattern with depth but depth was still associated with a significant amount of remaining variance. Primary productivity could not explain a significant amount of variation associated with longitude.

ANCOM-BC analyses were repeated to identify ASVs that differed across depth and longitude (**Supplementary File 2**). All major patterns were consistent including the depth trends of *Ca. Thalassoplasma*, *Endozoicomonas* ASVs, and *Shewanella* ASVs, as well as the site differences in corallicolids. The most notable exception was *Ca. Oceanoplasma* C1 which did not differ significantly with depth. 55 of 60, 32 of 45, and 52 of 77 ASVs that were significant in this manuscript were also significant in this older dataset between sites, along depth, and along longitude respectively.

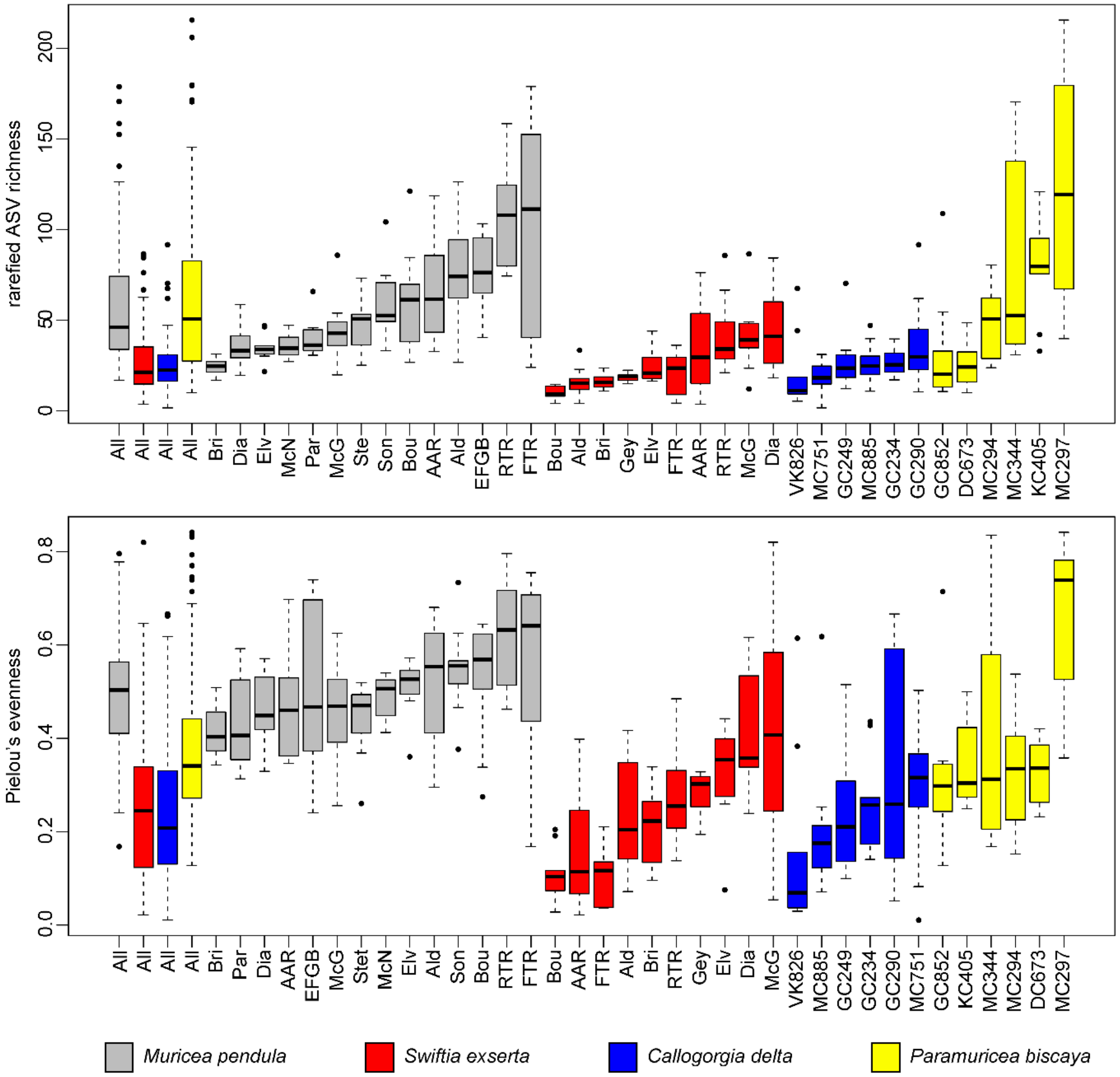


**Supplemental Figure 1.**

Differences in rarefied ASV richness and Pielou's evenness across sites and coral species. Boxplots are ordered by species and by sites in increasing order.

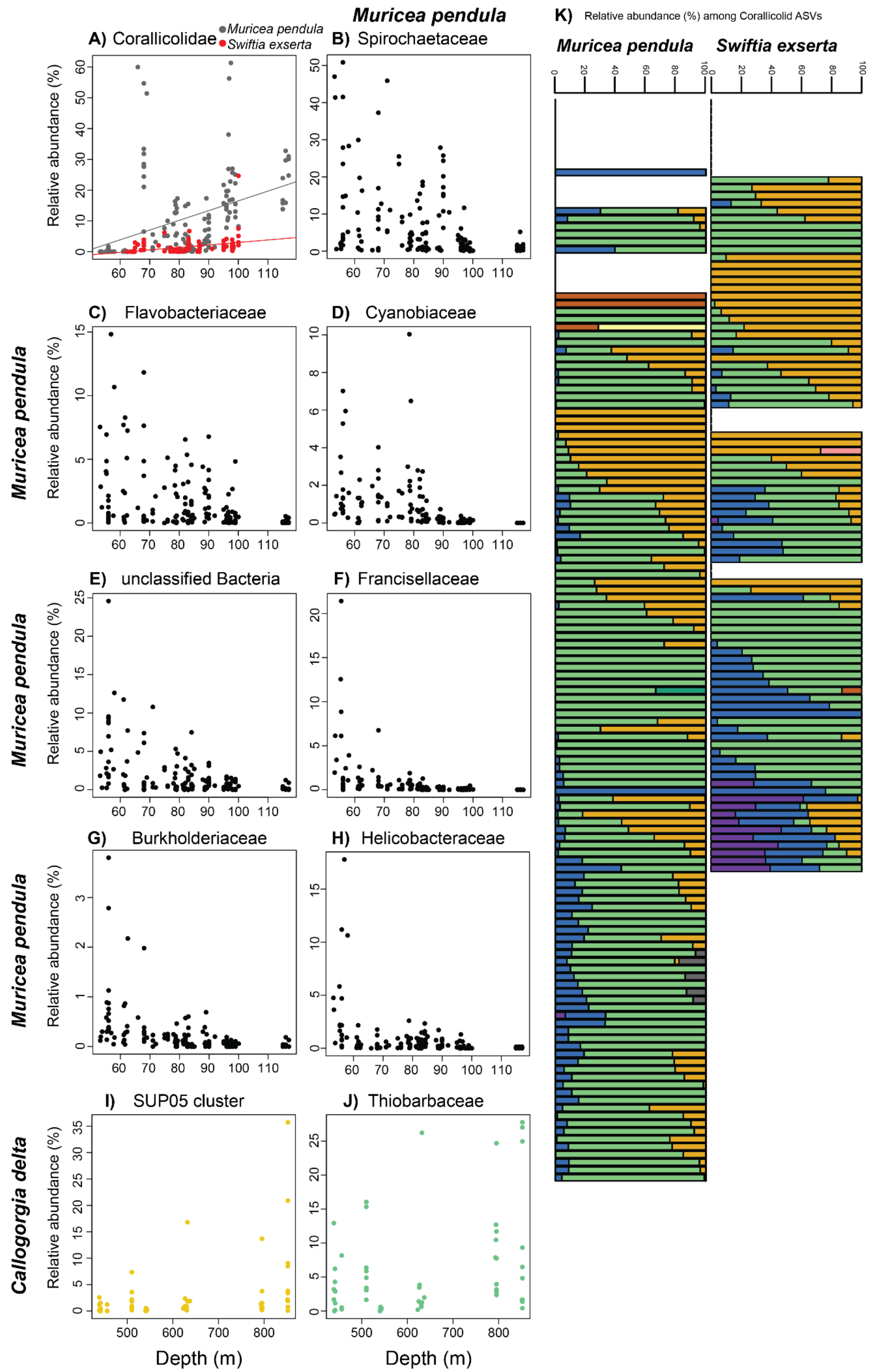

A) Corallicolidae
Muricea pendula
Swiftia exserta
Muricea pendula
B) Spirochaetaceae
K) Relative abundance (%) among Corallicolid ASVs
Muricea pendula
Swiftia exserta
C) Flavobacteriaceae
D) Cyanobiaceae
E) unclassified Bacteria
F) Francisellaceae
G) Burkholderiaceae
H) Helicobacteraceae
I) SUP05 cluster
J) Thiobarbaceae
Callogorgia delta
Relative abundance (%)
Depth (m)

**Supplemental Figure 2.**

More patterns with depth. A) Relative abundance of corallicolid ASVs in *M. pendula* (gray) and *S. exserta* (red) with depth. Relative abundance of the families B) Spirochaetaceae, C) Flavobacteriaceae, D) Cyanobiaceae, E) unclassified bacteria, F) Francisellaceae, G) Burkholderiaceae, H) Helicobacteraceae in *M. pendula* with depth. Relative abundance of I) SUP05 cluster and J) Thiobarbaceae in *C. delta* with depth. K) Relative abundance among corallicolid ASVs in *M. pendula* and *S. exserta* along depth. Each row is a coral sample and each color designates a different corallicolid ASV. Samples are grouped by site and sites are ordered by increasing depth.

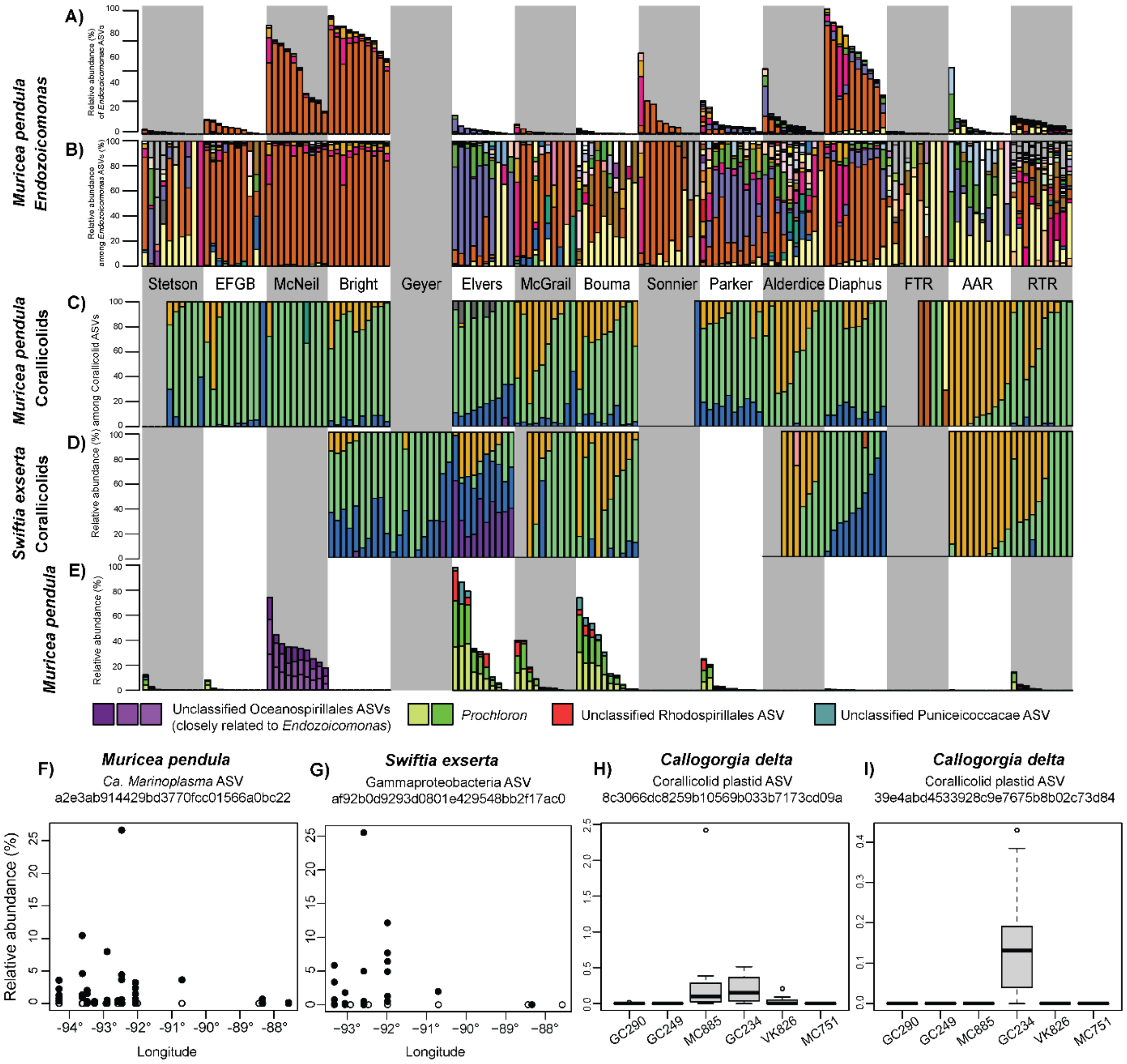


**Supplemental Figure 3**

Additional patterns with longitude and between sites. The relative abundance of *Endozoicomonas* ASVs A) and only among them B) in *M. pendula*. Samples are grouped by site and ordered from westernmost to easternmost. Relative abundances among corallicolid ASVs in C) *M. pendula* and D) *S. exserta*. E) Relative abundance of select ASVs that differ between sites in *M. pendula*. Relative abundance of F) a *Ca. Marinoplasma* ASV in *M. pendula* and G) an unclassified Gammaproteobacterial ASV in *Swiftia exserta* by longitude showing higher abundance in the west. Open circles denote samples with 0% relative abundance. H-I) Relative abundance of corallicolids by site in *C. delta*.

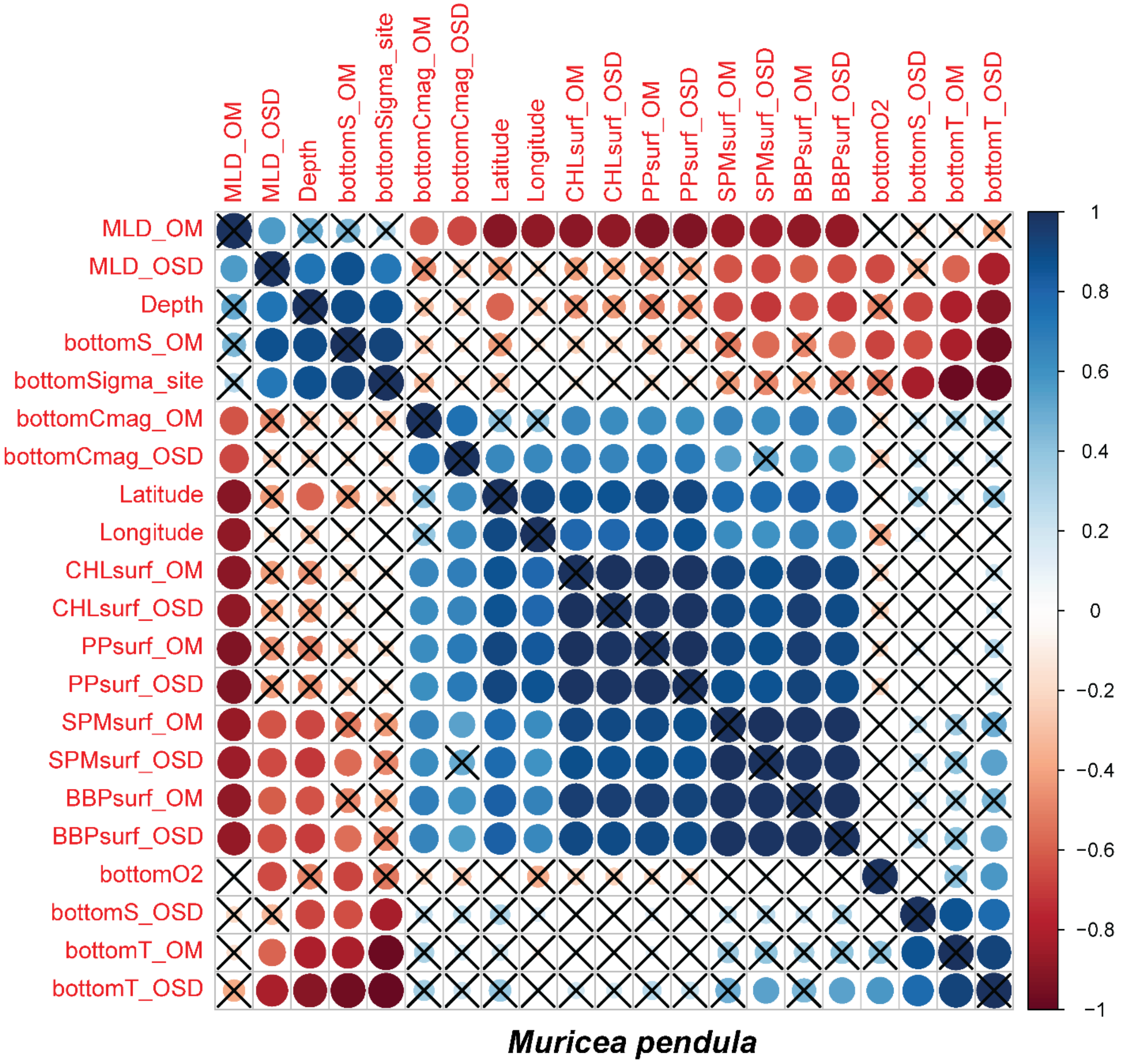


**Supplemental Figure 4.**

Correlation plot of environmental variables associated with *Muricea pendula* samples. Blue and red correspond to positive and negative correlations. X's mark variables that are not significantly correlated. OM: mean, OSD: standard deviation, surf: surface, MLD: depth of the mixed layer,

bottomS: salinity, bottomCmag: current magnitude, CHLsurf: chlorophyll concentration, SPM: suspended particulate matter, BBP: backscattering coefficient, bottomO2: oxygen concentration, bottom: temperature.

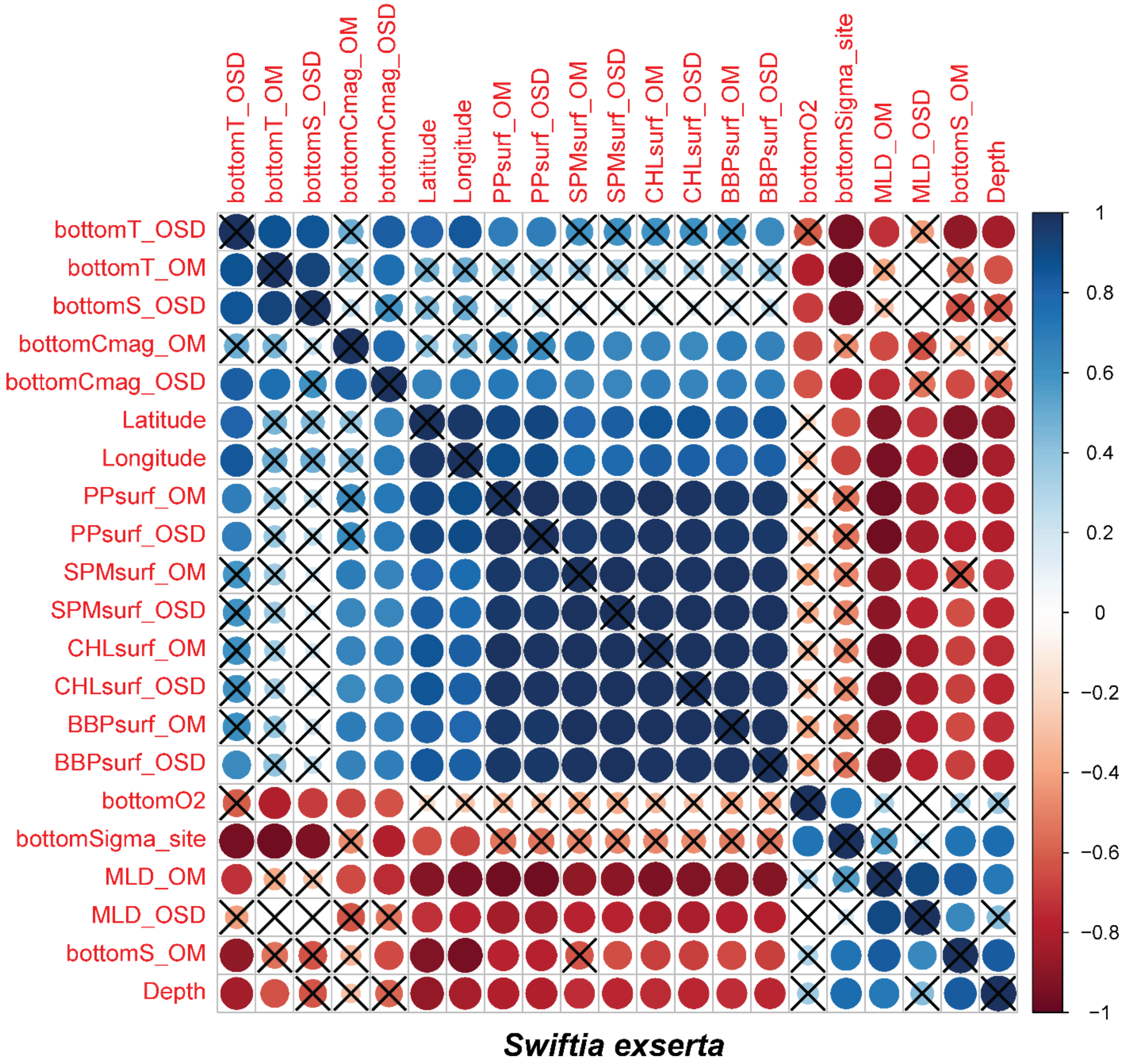


**Supplemental Figure 5.**

Correlation plot of environmental variables associated with *Swiftia exserta* samples. Blue and red correspond to positive and negative correlations. X's mark variables that are not significantly correlated. OM: mean, OSD: standard deviation, surf: surface, MLD: depth of the mixed layer,

bottomS: salinity, bottomCmag: current magnitude, CHLsurf: chlorophyll concentration, SPM: suspended particulate matter, BBP: backscattering coefficient, bottomO2: oxygen concentration, bottom: temperature.

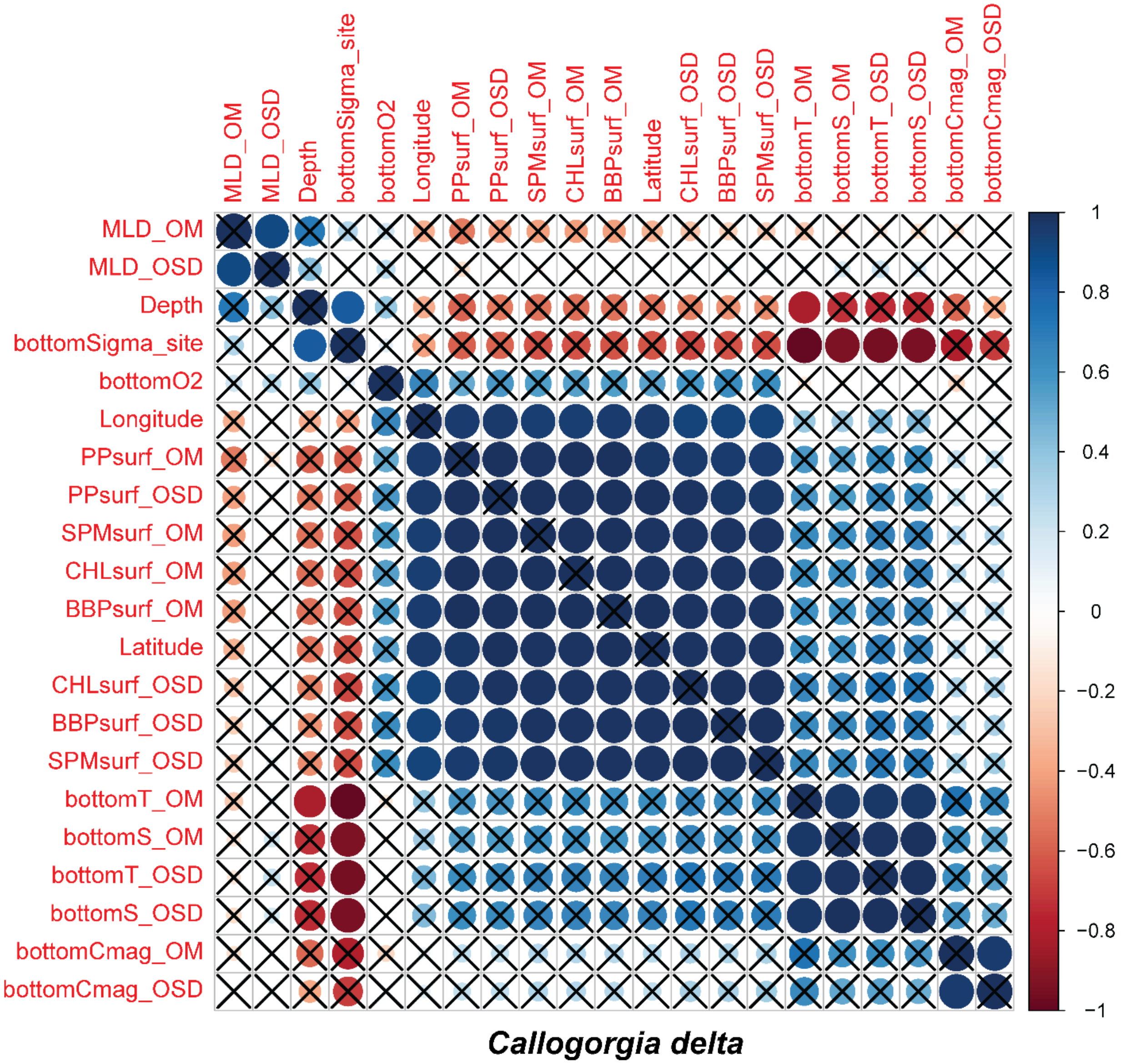


**Supplemental Figure 6.**

Correlation plot of environmental variables associated with *Callogorgia delta* samples. Blue and red correspond to positive and negative correlations. X's mark variables that are not significantly correlated. OM: mean, OSD: standard deviation, surf: surface, MLD: depth of the mixed layer,

bottomS: salinity, bottomCmag: current magnitude, CHLsurf: chlorophyll concentration, SPM: suspended particulate matter, BBP: backscattering coefficient, bottomO2: oxygen concentration, bottom: temperature.

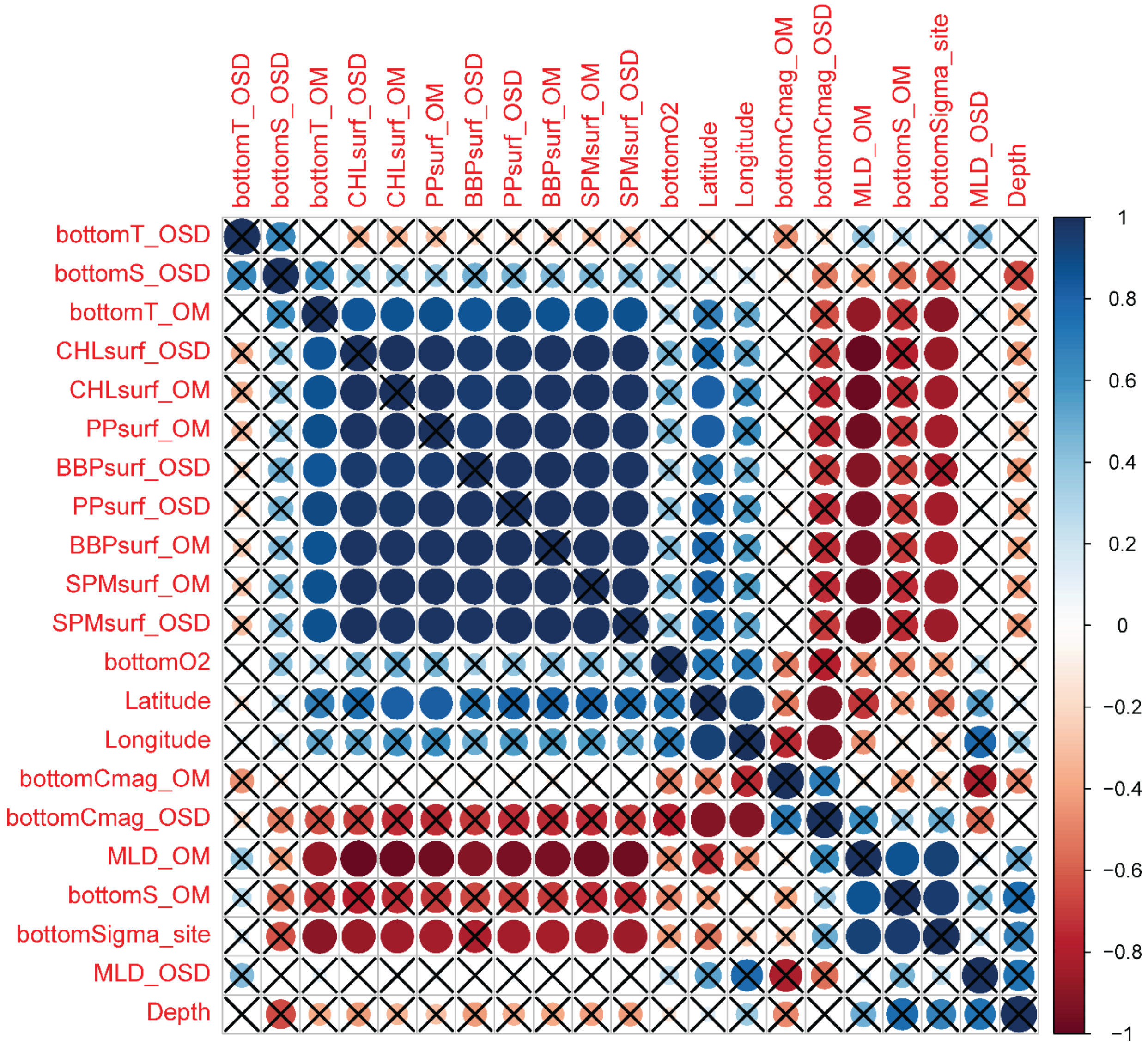


***Paramuricea biscaya***

**Supplemental Figure 7.**

Correlation plot of environmental variables associated with *Paramuricea biscaya* samples. Blue and red correspond to positive and negative correlations. X's mark variables that are not significantly correlated. OM: mean, OSD: standard deviation, surf: surface, MLD: depth of the mixed layer, bottomS: salinity, bottomCmag: current magnitude, CHLsurf: chlorophyll concentration, SPM: suspended particulate matter, BBP: backscattering coefficient, bottomO2: oxygen concentration, bottom: temperature.
